# Dietary amino acid manipulations in the common marmoset *Callithrix jacchus* impacts serum metabolites and FGF21 levels

**DOI:** 10.1101/2025.04.14.648667

**Authors:** Yuka Fujita, Tomoko Ishibuchi, Akiko Uematsu, Takuya Hayashi, Fumiaki Obata

## Abstract

Restricting dietary protein intake has metabolic and physiological benefits for animals. Rodent studies have identified the involvement of a hormone fibroblast growth factor 21 (FGF21), which is upregulated by sensing amino acid scarcity. However, to what extent this mechanism is conserved in primates remains elusive. Using common marmosets *Callithrix jacchus* as a non-human primate model, we demonstrate a protocol for protein restriction and dietary amino acid manipulation. Low protein diet induces a decrease in blood urea nitrogen, altered plasma amino acid profiles, and an increase in plasma FGF21. Supplementation of purified amino acids to the diet suppresses plasma features of protein restriction. Our data provide a dietary intervention technique in marmosets and an insight into the evolutionarily conserved mechanism of FGF21 induction during protein restriction.

## 1. Introduction

Dietary restriction is one of the most practical interventions for improving health. In addition to its benefits for patients with metabolic diseases^1^, there is mounting evidence that dietary restriction can enhance metabolic homeostasis and improve markers of longevity in healthy humans^2–4^, such as DNA methylation profiles^3^, anti-inflammatory responses in adipose tissue^2^, or cardio-metabolic parameters^4^. The composition of macronutrients plays a major role in the metabolic benefits of dietary restriction. For example, epidemiological studies have implied a correlation between mild restriction of dietary protein and metabolic benefits ^5,6^. Although limited in number, clinical interventional trials have demonstrated the potential physiological benefits of protein restriction on glucose homeostasis and lipid metabolism in healthy subjects^7,9,11^. Despite its potential impact, the mechanisms underlying these benefits of protein restriction in humans remain to be elucidated.

Molecular mechanisms of protein restriction have been extensively studies in rodents. In mice, dietary protein restriction has been reported to not only improve metabolic homeostasis, but also extend lifespan, independent of caloric intake^8,10^. With the use of genetics, fibroblast growth factor 21 (FGF21) has been highlighted as playing a major role in the improvement of glucose homeostasis and promotion of longevity associated with protein restriction^11,16,44^. FGF21 is a hormone synthesized and secreted from the liver, regulating the whole-body metabolism via its receptor expressed in various organs including liver, adipose tissue, and brain^13^. While FGF21 originally drew interests for its metabolic benefits^14^, its broader biological significance lies in its role as a key regulator that promotes adaptive metabolic stress response under imbalanced diet, characterized by increased food intake and energy expenditure, alongside suppressed growth and adiposity^14–18^. FGF21 is induced predominantly via activation of transcription factor 4 (ATF4) which is regulated by amino acid availability.

Interestingly, numerous studies have demonstrated that the deficiency of select few, in some cases single, amino acids can induce FGF21 in rodents^14,15,17–19^. While many amino acids are decreased in the liver and plasma during protein restriction, it has been reported that addition of just threonine and tryptophan to the low protein diet repressed the FGF21 induction^19–21^. The deficiency in dietary threonine or tryptophan was also reported to be sufficient for the induction of FGF21. These studies suggest that not all amino acids contribute equally to the induction of FGF21 in protein restriction, with threonine and tryptophan playing particularly critical roles. According to clinical studies, the FGF21 induction in response to protein restriction is also conserved in humans^7,9,11^. However, the molecular mechanisms and the amino acid profiles underlying this induction in primates, including humans, remain elusive.

To promote translational studies, primate models has been employed in dietary intervention studies^23–25^. Oral administration of FGF21 analogue is effective for weight loss and improvement of glucose homeostasis in various primate metabolic disease models ^26^. However, whether and how protein restriction induces endogenous FGF21 in non-human primate models has not been studied. Identifying the precise role of individual amino acids in FGF21 induction in primates requires the use of a well-defined diet with an easily adjustable amino acid composition. The high cost of purified amino acids required for dietary manipulation presents a challenge, particularly in primates that are generally much larger in size than rodents. To address this, we employed a smaller primate, common marmoset *Callithrix jacchus*. Common marmosets are new world monkeys endemic to South America, and are increasingly being used as a primate model for various biochemical research due to their small size (approximately 250-500 g for adults^27^) and the relative ease of breeding^26^. While marmosets are especially utilized in the field of neuroscience, they are also becoming a model to study infectious diseases, aging, and metabolic diseases. They are omnivores, with natural diet consisting of insects, fruits, and tree exudates such as sap and gum^28^. In captivity, marmosets are fed varying commercially available diets enriched with different snacks. While challenges remain in the optimizing captive diets to minimize individual variations in digestion^29^, marmosets are an optimal model to translate the findings from rodent studies to humans in the field of nutritional research. In this study, we established dietary manipulation of protein and amino acids in marmosets, and investigated the plasma profiles of BUN, amino acids, and FGF21.

## 2. Results

### Determination of standard diet for dietary manipulation

To manipulate dietary composition, we decided to define the standard diet for common marmosets. First, we surveyed the current state of marmoset diet in captivity to determine the macronutrient composition. Commercially available diets for primates include Oriental SPS and CMS-1M from two leading manufacturers for animal diets in Japan. We also investigated the marmoset diets used in a Japanese zoo to estimate weekly averaged intake of nutrient. In all the cases, the macronutrient in the diet was made up of 20-30% protein (Supplementary Figure 1a). Based on this diet, we determined a control diet named P30, consisting of 30 % protein in calories (Table 1).

**Table 1.** Dietary composition of the experimental diet.

|  |  |  |  |  |
| --- | --- | --- | --- | --- |
|  | P30 diet<br>(D24021601M) |  | P5 diet<br>(D24021602M) |  |
|  | gm | kcal | gm | kcal |
| Protein (%) | 30.5 | 30 | 5.1 | 5 |
| Carbohydrate (%) | 48.8 | 48 | 74.3 | 73 |
| Fat (%) | 10 | 22 | 10 | 22 |
| Total |  | 100 |  | 100 |
| kcal/gm | 4.08 |  | 4.08 |  |
| Ingredient | gm | kcal | gm | kcal |
| Casein | 300 | 1200 | 50 | 200 |
| L-Cystine | 4 | 16 | 0.7 | 3 |
| Corn Starch | 320 | 1280 | 573.3 | 2293 |
| Maltodextrin 10 | 100 | 400 | 100 | 400 |
| Sucrose | 65 | 260 | 65 | 260 |
| Cellulose, BW200 | 50 | 0 | 50 | 0 |
| Soybean | 100 | 900 | 100 | 900 |
| Mineral Mix,<br>S40001 | 50 | 0 | 50 | 0 |
| Vitamin Mix,<br>V40001C | 1 | 4 | 1 | 4 |
| Choline Bitartrate | 2 | 0 | 2 | 0 |
| L-Ascorbic Acid,<br>Phosphate | 3 | 0 | 3 | 0 |
| (33% Ascorbic<br>acid) |  |  |  |  |
| Vitamin D3,<br>100,000 IU/gm | 3.283 | 0 | 3.283 | 0 |
| Total | 995.279 | 4060 | 995.279 | 4060 |

Many biomarkers have been proposed to reflect the adequate protein intake in humans^30,31^. Nitrogen, especially blood urea nitrogen (BUN) is used to monitor internal amino acid availability^32–34^. Plasma BUN levels from marmosets fed the P30 diet for two weeks showed no significant alteration than the conventional CMS-1M-based diet, indicating that the P30 diet contains sufficient protein content (Supplementary Figure 1b).

### Protein restricted diet decreases BUN and induces FGF21 in plasma

For protein restriction, we designed an isocaloric low protein diet, P5 which contains 5% protein (Supplementary Fig 1A, Table 1) and characterized the metabolic responses of male marmosets to this diet (Figure 1A). Marmosets on the low protein diet did not show significant changes in body weight (Figure 1B), or plasma albumin levels (Figure 1C). In contrast, BUN levels strongly decreased during protein restriction (Figure 1D).

**Figure 1.**
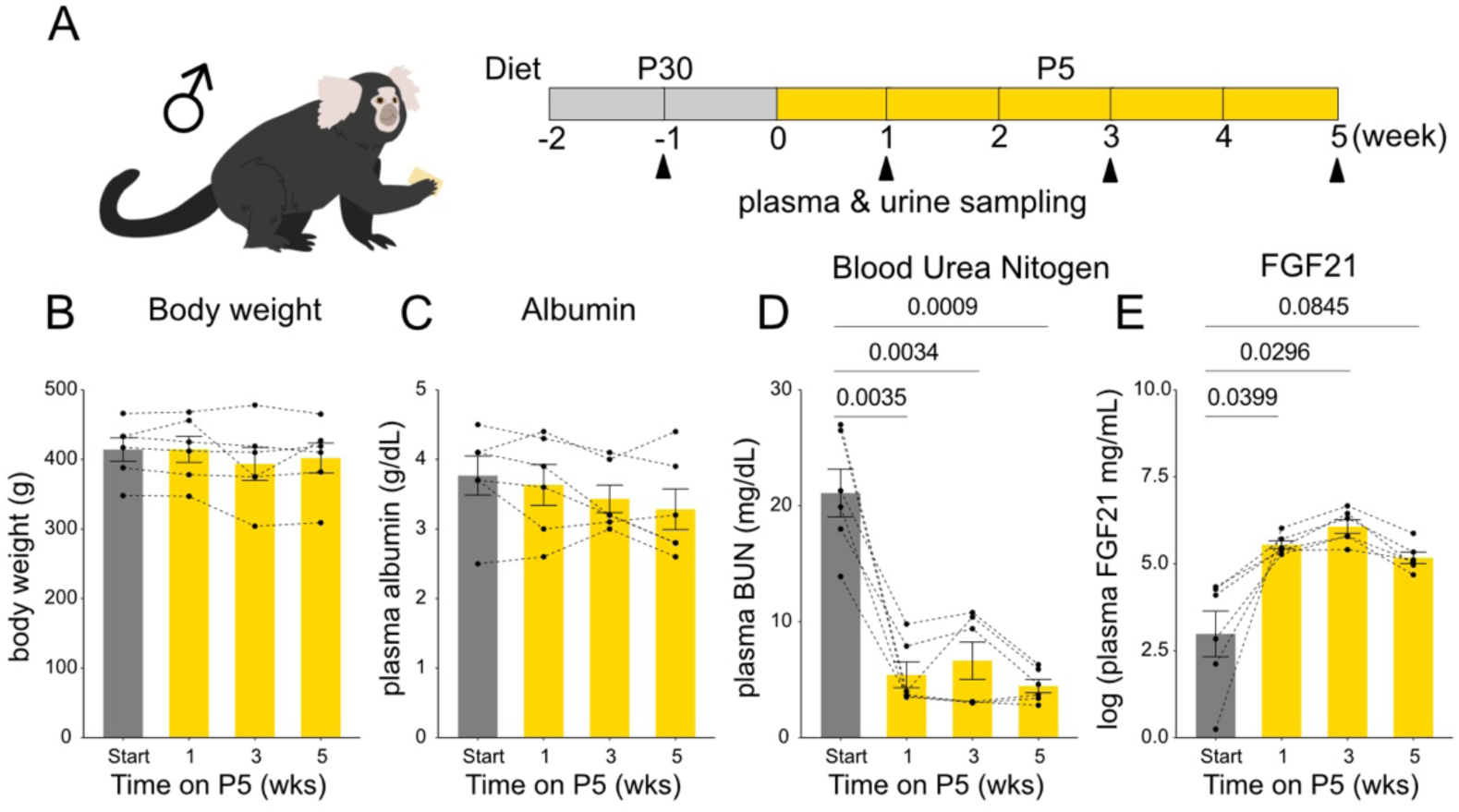
Plasma markers of low protein is observed by P5 diet (A) Experimental timeline. Six male marmosets were fed P30 diet for two weeks and P5 diet for 5 weeks. Blood and urine samples were obtained every two weeks. (B-E) Bodyweight (B), plasma albumin (C), and plasma blood urea nitrogen (BUN) (D), and plasma FGF21(E) levels. For (B-E), statistical analysis was performed by repeated measures one-way analysis of variance (ANOVA). The *p* values determined by post hoc analysis with Dunnet’s multiple comparison test are shown in the figure. All data are presented as mean ± s.e.m.

As expected, induction of FGF21 occurred at as early as one week on the low protein diet (Figure 1E). These data suggest that one week on a 5% protein diet was sufficient to decrease BUN and induce FGF21, demonstrating typical responses to protein restriction found in rodents.

### Plasma metabolic profile of marmosets is altered on low protein diet

To understand how the low protein diet affected levels of internal amino acids and the related metabolites, we quantified amino acids and taurine in the plasma using LC-MS/MS. Cysteine was excluded from the analysis due to technical issues in reliable measurement. Taurine was quantified since it is an amino acid metabolite which is known to be highly influenced by dietary amino acids^34,35^. Principle component analysis (PCA) of the quantified 20 metabolites showed that protein restriction separates the plasma metabolite profile (Figure 2A). We found that essential amino acid levels especially responded to the P5 diet (Figure 2B-D, Supplementary Figure 2). Among essential amino acids, threonine (Figure 2C) and tryptophan (Figure 2D) showed significant drops in plasma concentration after 1 week of protein restriction. Plasma concentration of several non-essential amino acids were rather increased on the P5 diet (Figure 2E-F, Supplementary Figure 2). Although p=0.0502 in repeated measures one way ANOVA, we observed a trend of increase in alanine concentration, suggesting potential differences. Dunnet’s post hoc analysis demonstrated that alanine indeed tended to increase under protein restriction (Figure 2F). Surprisingly, methionine, despite being an essential amino acid, increased after two weeks of protein restriction (Figure 2G). Upon closer inspection, we instead found that taurine, which is synthesized from methionine, was significantly decreased under protein restriction (Figure 2H-I). Consistent with plasma, we found that urinary taurine was also decreased by the low protein diet (Figure 2J). Taken together, our data showed that while one week of P5 diet is sufficient to induce the adaptative mechanism towards protein restriction, only selected amino acids decrease in the plasma.

**Figure 2.**
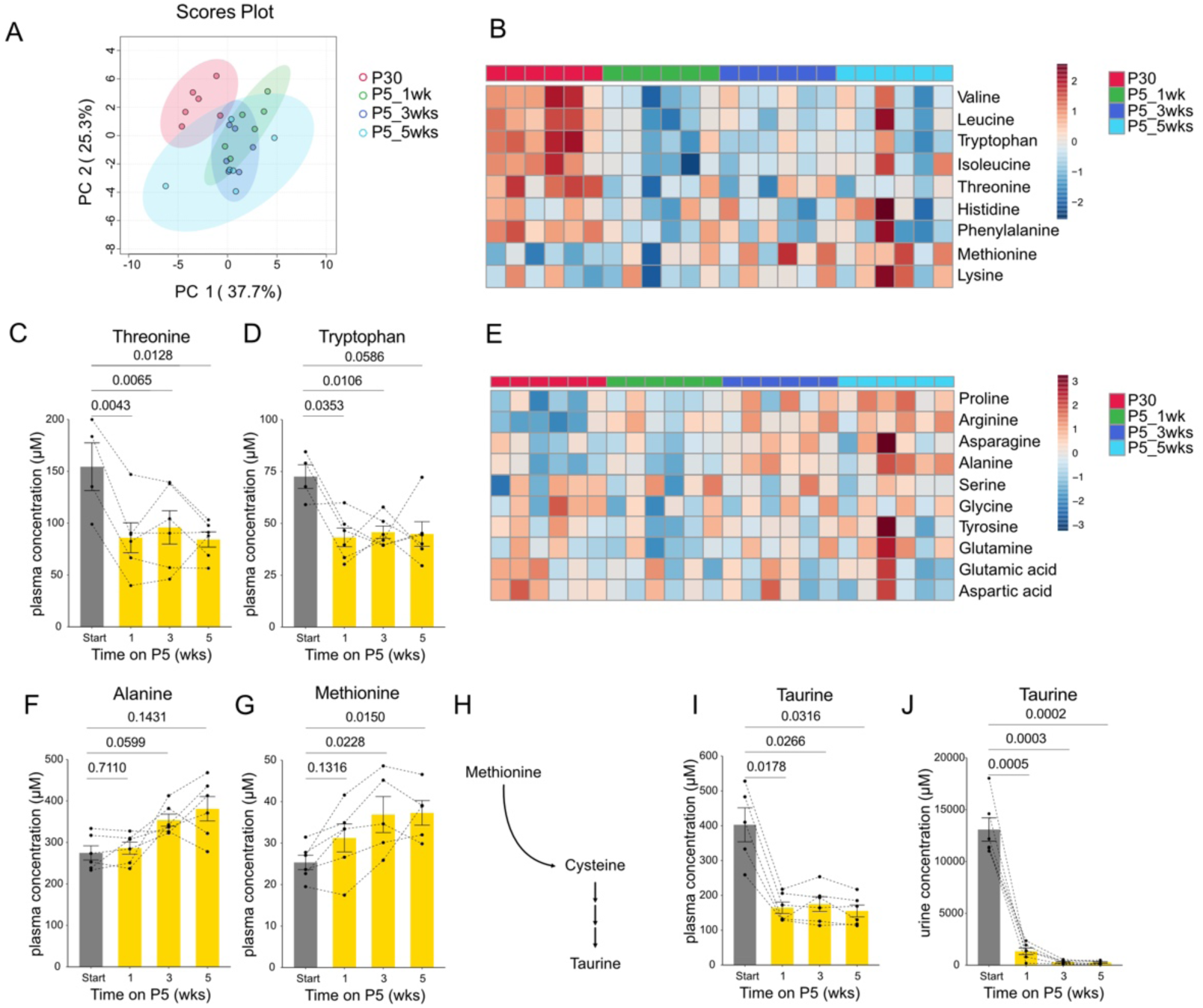
Plasma metabolite profiles are altered under P5 diet (A) Principal component analysis of plasma metabolites in male marmosets quantified using LC-MS/MS. (B) Heatmap of plasma essential amino acids. (C-D) Quantification of plasma threonine (C) and tryptophan (D). (E) Heatmap of plasma non-essential amino acids. (F-G) Quantification of plasma alanine (F) and methionine (G). (H) Schematics of taurine biosynthesis pathway. (I-J) Quantification of taurine in plasma (I) and urine (J). For (C-D, F-G, I-J), statistical analysis was performed by repeated measures one-way analysis of variance (ANOVA). The *p* values determined by post hoc analysis with Dunnet’s multiple comparison test are shown in the figure. All data are presented as mean ± s.e.m.

### Decreased BUN and FGF21 during protein restriction is ameliorated upon free amino acid supplementation

To test whether these responses to the low protein diet is owing to a decrease in amino acid levels and, if so, to determine its threshold, we fed the animals with diets containing different levels of free amino acids. The composition of amino acids was matched to casein (Figure 3A, See Methods, Table 2-4). P30, P5, and P5 diet supplemented with free amino acids, were administered sequentially for one week each (Figure 3B). Since the diets were provided *ad libtum,* we deemed their isocaloric status less relevant, and thus we additionally supplemented free amino acids to a set amount of P5 food (Figure 3A). The tested diets included P30 diet, P5 diet, and P5 diet supplemented with free amino acid cocktail corresponding to 5%, 15%, or 25% of casein (Figure 3A). More specifically, amino acids were supplemented at an amount identical to (P5+5%), three times (P5+15%), or five times (P5+25%) the predicted content in the P5 diet. P5+25% diet is supposed to contain the same amount of amino acids as P30.

**Figure 3.**
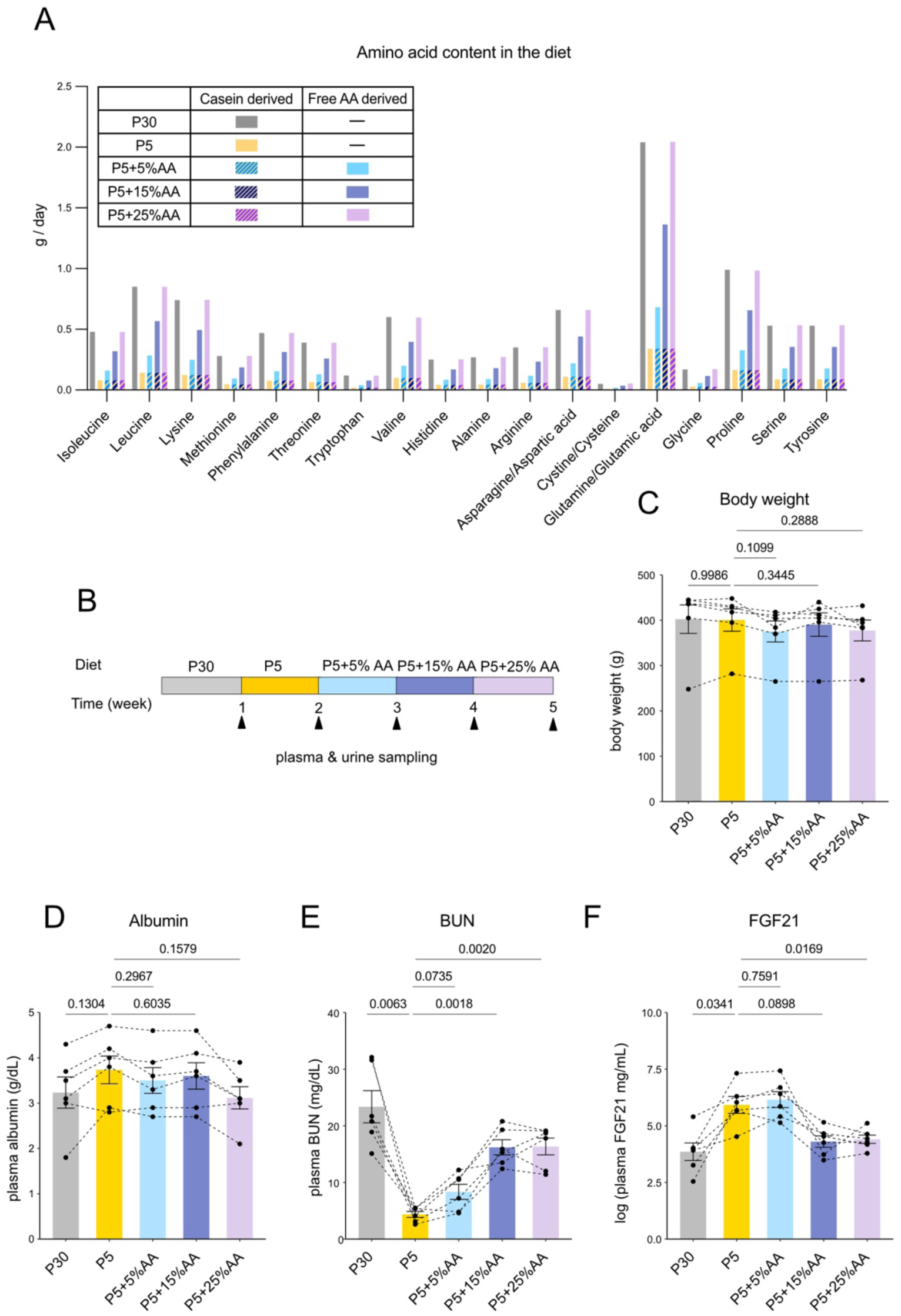
Decrease in BUN and increase in FGF21 levels upon protein restriction is repressed by amino acid supplementation (A) Amount of amino acid derived from casein or reagent supplementation contained in each diet by weight per day per marmoset. 30g of food was supplied daily per subject. (B) Experimental timeline. Six male marmosets were fed P30, P5, and P5 diet supplemented with free amino acids for 1 week each. Blood and urine samples were obtained weekly. The diet was fed *ad libtum.* (C-F) Bodyweight (C), plasma albumin (D), and plasma blood urea nitrogen (BUN) (E), and plasma FGF21 levels (F). For (C-F), statistical analysis was performed by repeated measures one-way analysis of variance (ANOVA). The *p* values determined by post hoc analysis with Šídák’s multiple comparison test are shown in the figure. All data are presented as mean ± s.e.m.

**Table 2.** Amino acid composition of casein.

| Amino Acid Profile | (g/100g of casein protein) |
| --- | --- |
| Isoleucine | 5.3 |
| Leucine | 9.4 |
| Lysine | 8.2 |
| Methionine | 3.1 |
| Phenylalanine | 5.2 |
| Threonine | 4.3 |
| Tryptophan | 1.3 |
| Valine | 6.6 |
| Histidine | 2.8 |
| Alanine | 3 |
| Arginine | 3.9 |
| Aspartic acid | 7.3 |
| Cysteine/Cystine | 0.6 |
| Glutamic acid | 22.6 |
| Glycine | 1.9 |
| Proline | 10.9 |
| Serine | 5.9 |
| Tyrosine | 5.9 |

Throughout the serial dietary manipulation, health parameters such as body weight, or plasma albumin levels did not vary significantly (Figure 3C, D). However, the decreased BUN levels under P5 diet gradually increased with amino acid supplementation (Figure 3E). Importantly, the same pattern was also observed in the plasma concentration of FGF21 (Figure 3F). The induction upon protein restriction by P5 was not changed with P5+5% diet but was repressed at P5+15%. Considering that P5+15% almost seemed to completely abrogate the decrease of BUN and induction of FGF21, and that further increase of supplemented free amino acids could not influence them, we concluded that P5+15% was sufficient for the observed responses (Figure 3E-F). We also noticed that BUN had a trend to increase in P5 + 5%, whereas FGF21 did not, suggesting that BUN acts more sensitively to reflect dietary protein or amino acid levels whereas the FGF21 induction has a clear threshold that responds to the low amino acid status.

### Plasma metabolite levels upon free amino acid supplementation

Plasma samples were analyzed using LC-MS/MS to assess metabolite profiles under amino acid supplementation. PCA showed that amino acid supplementation progressively restored the plasma metabolomic profile in accordance with increasing amino acid content of protein restriction, mainly along PC3 (Figure 4A).

**Figure 4.**
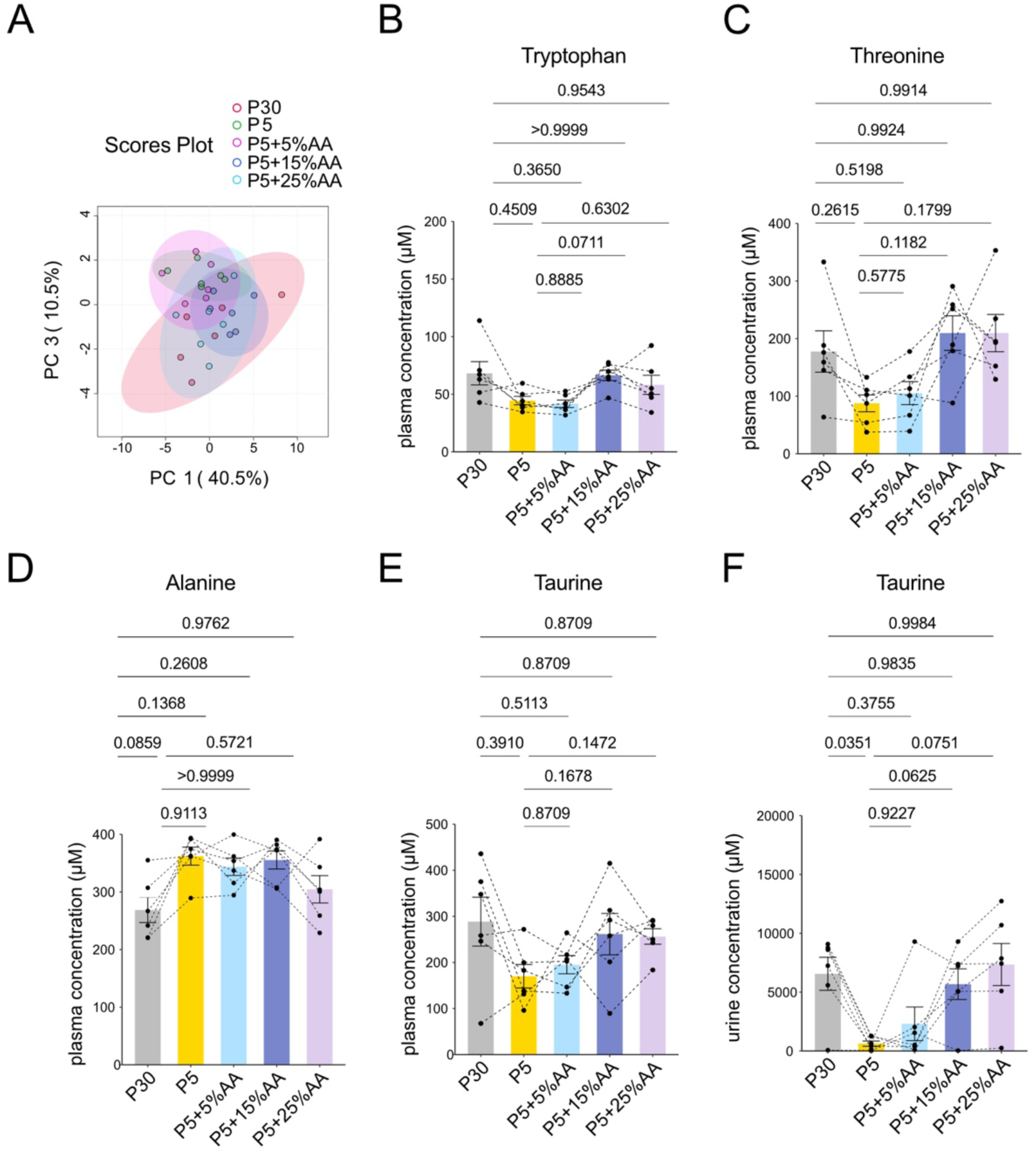
Amino acid supplementation can ameliorate the metabolite profile of low protein diet (A) Principal component analysis of plasma metabolites in male marmosets quantified using LC-MS/MS. (B-D) Quantification of plasma tryptophan (B), threonine (C), alanine (D). (E-F) Quantification of taurine concentration in plasma (E) and urine (F). For (B-F), statistical analysis was performed by repeated measures one-way analysis of variance (ANOVA). The *p* values determined by post hoc analysis with Šídák’s multiple comparison test are shown in the figure. All data are presented as mean ± s.e.m.

Tryptophan, threonine and alanine were the only amino acids with variable change (Figure 4B-D, Supplementary Figure 3). As observed in Fig. 2, threonine and tryptophan tended to decrease, and alanine tended to increase on the P5 diet, although the change was not statistically significant (Figure 4B-D). For tryptophan and threonine, plasma concentrations during P5+15%AA diet tended to increase the most compared to the level observed in P5 (Figure 4B-C). In contrast, alanine showed significant increase by the P5 diet and its levels were at least partially restored by P5+25%AA diet (Figure 4D). In this experiment, methionine levels were unchanged after one week on the P5 diet (Supplementary Figure 3). Mean plasma taurine levels showed a decreasing trend following one week on P5 diet. Consistent with previous results (Figure 2J), urinary taurine also decreased on P5, and this reduction tended to be suppressed by amino acid supplementation (Figure 4F).

## Discussion

In this study, we established the experimental set up for dietary protein and amino acid manipulation in common marmosets. The nutritional responses of marmosets to a low protein diet included a decrease in BUN, an induction of FGF21, and altered plasma amino acid levels that occurred within a week.

Amino acids that especially decreased under low protein diet were threonine and tryptophan. This result was consistent with previous studies using rodents^35,36^. The reasons that threonine and tryptophan seem to be the most sensitive to low protein diet is unknown, but the two amino acids have distinct characteristics that may contribute to the sensitivity to low protein diet. For example, threonine is known to be especially utilized in the gut and thus is reported to have high irreversible ileal loss in humans^37^. One major factor is secretion of glycoprotein, such as mucin, to the lumen of the gut.

Mucin is extensively O-glycosylated, requiring an extremely high proportion of amino acids with hydroxylated side chains, mainly threonine and serine^36^. Since mucin is secreted into the intestinal lumen and cannot be reutilized due to its low digestibility and low absorptive abilities in the colon, a significant portion of threonine is predicted to be lost. Total loss of threonine at the terminal ileum of healthy humans has been calculated to account for 103-127% of total daily maintenance requirement^15,38^. This is significantly higher compared to the ileal losses of other amino acids estimated to be 26% (Histidine)-75%(Valine). Previous researches in mice have also discussed that threonine may be the limiting amino acid in a casein diet^39^.

On the other hand, tryptophan is structurally distinct, being the only amino acid with an indole ring. It has diverse metabolites from the kynurenine pathway to serotonin production. It is also highly metabolized by the gut bacteria, received mainly by the aryl hydrocarbon receptor in the host^40^. In addition, tryptophan, along with cystine is predicted to be the least abundant amino acid in human foods^32^. Amino acid with the lowest content in casein used in this study was tryptophan, at 1.3% (w/w) (Table 3). These characteristics may have led to the high likelihood of threonine and tryptophan acting as a rate limiting amino acid under low protein diet. However, whether this is the case still needs further to be investigated by manipulation of the two amino acids and tracking of the dynamics and distribution of amino acids from both endogenous and dietary sources. Relatedly, it is necessary to measure amino acids in the liver, where FGF21 is produced, since we have only measured amino acid levels in the blood in this study.

**Table 3.**
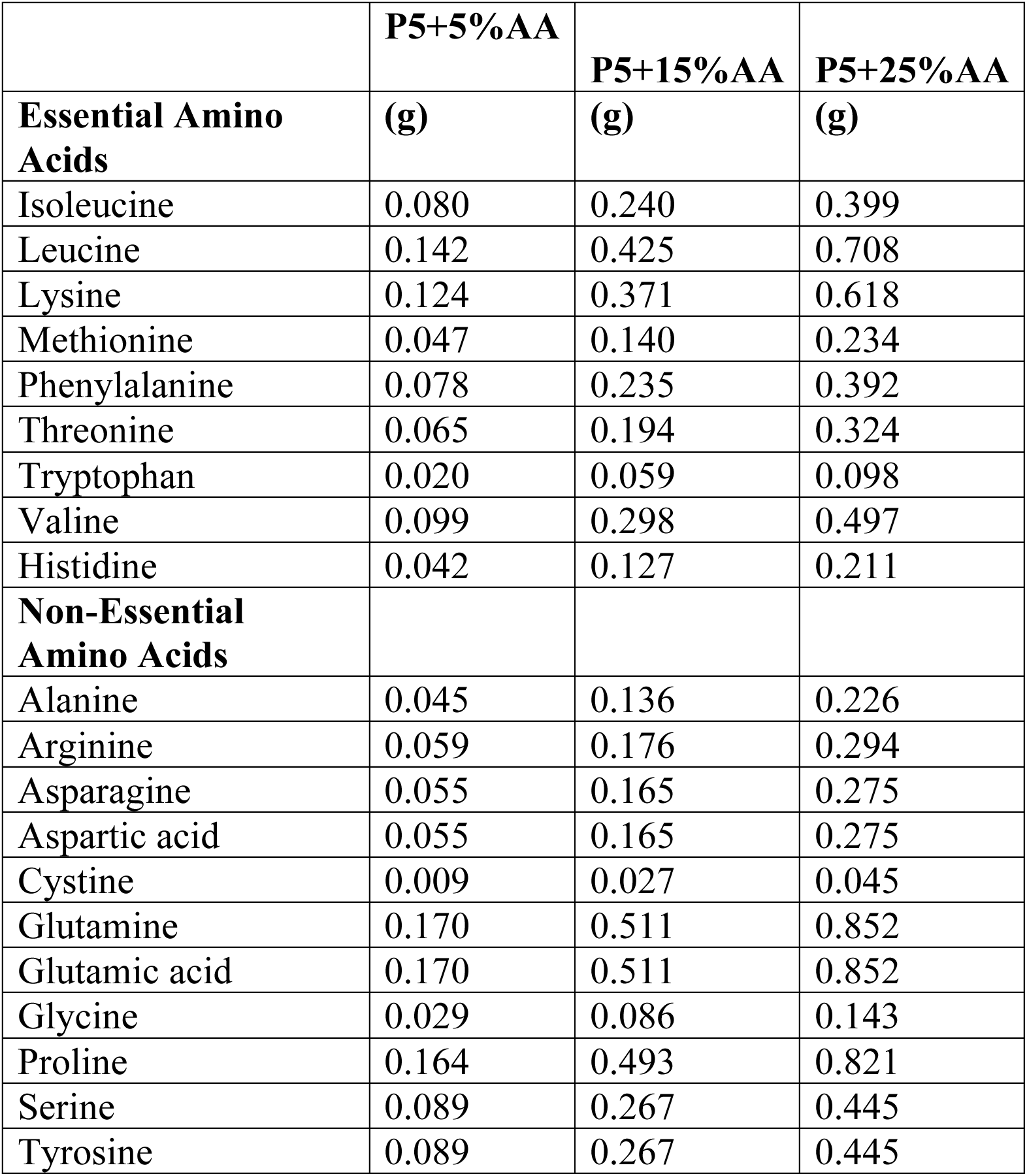
Free amino acids supplemented to 30 g of P5 diet.

|  | **P5+5%AA** | **P5+15%AA** | **P5+25%AA** |
| --- | --- | --- | --- |
| **Essential Amino Acids** | **(g)** | **(g)** | **(g)** |
| Isoleucine | 0.080 | 0.240 | 0.399 |
| Leucine | 0.142 | 0.425 | 0.708 |
| Lysine | 0.124 | 0.371 | 0.618 |
| Methionine | 0.047 | 0.140 | 0.234 |
| Phenylalanine | 0.078 | 0.235 | 0.392 |
| Threonine | 0.065 | 0.194 | 0.324 |
| Tryptophan | 0.020 | 0.059 | 0.098 |
| Valine | 0.099 | 0.298 | 0.497 |
| Histidine | 0.042 | 0.127 | 0.211 |
| **Non-Essential Amino Acids** |  |  |  |
| Alanine | 0.045 | 0.136 | 0.226 |
| Arginine | 0.059 | 0.176 | 0.294 |
| Asparagine | 0.055 | 0.165 | 0.275 |
| Aspartic acid | 0.055 | 0.165 | 0.275 |
| Cystine | 0.009 | 0.027 | 0.045 |
| Glutamine | 0.170 | 0.511 | 0.852 |
| Glutamic acid | 0.170 | 0.511 | 0.852 |
| Glycine | 0.029 | 0.086 | 0.143 |
| Proline | 0.164 | 0.493 | 0.821 |
| Serine | 0.089 | 0.267 | 0.445 |
| Tyrosine | 0.089 | 0.267 | 0.445 |

Taurine levels change drastically, especially in urine, in response to a low protein diet. Taurine is regulated by both biosynthesis from methionine or cysteine and renal reabsorption^41^. Additionally, taurine is known to be abundantly contained in the skeletal muscle which functions as an amino acid reservoir^32–34^. Restriction of dietary protein, or even just methionine has been reported to decrease plasma and hepatic taurine levels and alter its metabolism in rodents^42^. Interestingly, our results demonstrated that plasma methionine levels did not decrease under protein restriction, perhaps at the expense of taurine. Similarly, upon protein restriction, the levels of non-essential amino acids are well maintained, or can even increase, instead. In a previous study, plasma of male rats on protein restricted diet for four days showed an increase in serine, glycine, glutamine, glutamate, and alanine levels in the plasma^43^. In another study, maintaining female rats on low protein diet for 7-10 days induced an increase in hepatic serine and glycine^44,45^. The exact mechanism behind the upregulation is not fully understood. One possibility is the shift in metabolism, such as the induction of amino acid biosynthesis. A prior study using mouse demonstrated that transcriptional upregulation of *asparagine synthetase* (*ASNS*) and *phosphoglycerate dehydrogenase* (*PHGDH*), which are involved in asparagine and serine synthesis respectively, occurs under protein restriction via ATF4^46^. Decreased global translation and selective upregulation of the translation of proteins involved in adaptive responses under protein limitation has also been elucidated^47^. Taken together, a robust adaptive system is likely preventing a decrease in circulating amino acid levels by regulating absorption, excretion, and metabolism under dietary amino acid deficiency.

In our study, we set the control diet to 30% protein. In primate studies, the standard diet varies greatly among institutions^20–22,48^. The lack of consensus in defining standard diet have made it difficult to interpret the results of dietary manipulation among studies. For example, three well-known primate studies from different institutes which looked at the effect of calorie restriction reported varying degrees of positive effects on health and age-related morbidity, and inconsistent outcomes for all cause mortality^48,49^. The discrepancy in the efficacy of caloric restriction has been attributed to the diversity not only in animals’ age or genetic and metabolic background, but experimental design, such as dietary composition or feeding practices^50^. For the promotion of nutritional study, it is necessary to define the macronutrient composition of a standardized diet carefully.

In addition to macronutrient composition, the effects of individual nutrients should be considered when interpreting experimental results. Our study implied that individual amino acids may affect nutrient sensing pathways in marmosets, similarly to rodents.

Experimental animals in captivity are fed different sources of protein, mainly casein, whey, and soy which all have varied amino acid compositions that can influence experimental interpretation. For example, a previous study by Tardif *et al.* has reported that 15 or 25% protein diet did not alter growth and food intake behavior of infants^51^ nor milk macronutrient compositions collected from marmoset mothers ^52^. Consistently, we observed that 5% protein diet supplemented with free amino acids equivalent to 15%, but not 5% protein was sufficient to diminish the responses of protein restriction under the conditions tested. These results imply the possibility approximately 15% protein is sufficient to not induce phenotypes of low protein. However, such conclusion must be made cautiously, since the protein source for the experiments by Tardif et al was lactalbumin which notably contains higher amount of essential amino acids including isoleucine, leucine, and tryptophan compared to casein^53^. Definitive conclusions require further investigation with matched amino acid compositions and endocrine profiling. These comparisons shed light on the importance of manipulating specific dietary components at the amino acid levels, by using approaches like the one we proposed in this study.

## Limitations

The animals were fed *ad libtum*, meaning that the intake was completely dependent on the animals themselves. While we did not observe clear difference in appetite between the diets, we cannot be sure since the dietary intake was not quantified. Only matured male marmosets were available to test and there might be sex and life-stage differences.

## 3. Material and methods

### Animals

All experiments were performed in strict accordance with the institutional guidelines for animal experiments, basic policies for the conduct of animals experiments in research institution (MEXT, Japan), and guidelines for the care and use of laboratory animals (National Institute of Health, Bethesda, MD). All animal experiments were approved by the Animal Care and Use committee of the RIKEN Center for Biosystems Dynamics Research (Kobe, Japan) (approved number: A2023-27-2).

Subjects were male marmosets (*Callithrix jacchus*) housed in RIKEN BDR. The monkeys were born at the center and maintained in an individual or family style cage before the experiment. The subjects were 6.7±0.94 years of age at the start of the experiments. During the experiments, the monkeys were and individually housed in a cage (450 × 660 × 600 mm) with a wooden perch for enrichment. The cages were placed facing each other, so subjects were exposed to visual, auditory and olfactory cues from each other. The room was maintained at 28 degrees and 50% humidity, with 12 hours light-dark cycle (Light: 8:45-20:45).

Diets were given twice a day and food supplements were given three times a week, approximately 5g of banana or 3g of boiled eggs, twice and once a week respectively throughout the experiment. Prior to the start of the experiment, the diet consisted of commercial New World primate diet (CMS-1M, CLEA Japan) supplemented with honey (Kato brothers honey), ascorbic acid (Kenei), vitamin mix (Vitalong, Kyoritsu Seiyaku) and probiotics (BIO-THREE, TOA).

### Experimental Diet and feeding

Two types of experimental diets, P30 (D24021601M) and P5 (D24021602M), were purchased from Research Diets (NJ, USA) in powder form. The composition of each diet can be found in Table1. The amount of individual amino acids predicted to be in the diet containing 5% protein, termed P5 diet was calculated based on the composition of casein provided by Research Diets (Table 2). P5+5%AA, P5+15%AA, and P5+25%AA diet were prepared by mixing a set amount of P5 diet with 100%, 300%, or 500% of amino acids predicted to be in the set amount of P5 diet (Table 1-3). Amino acid reagents used in this study can be found in Table 4. The powder diet was mixed with 7.2 mL of honey diluted with 70 mL (80 mL for P5 diet) of water, 1 g of probiotics (BIO-THREE, TOA) and left to set in a metal tray. The diet was then cut into bite sized pieces before feeding. The diet was prepared up to 3 days in advance and kept in a fridge at roughly 4 degrees.

**Table 4.**
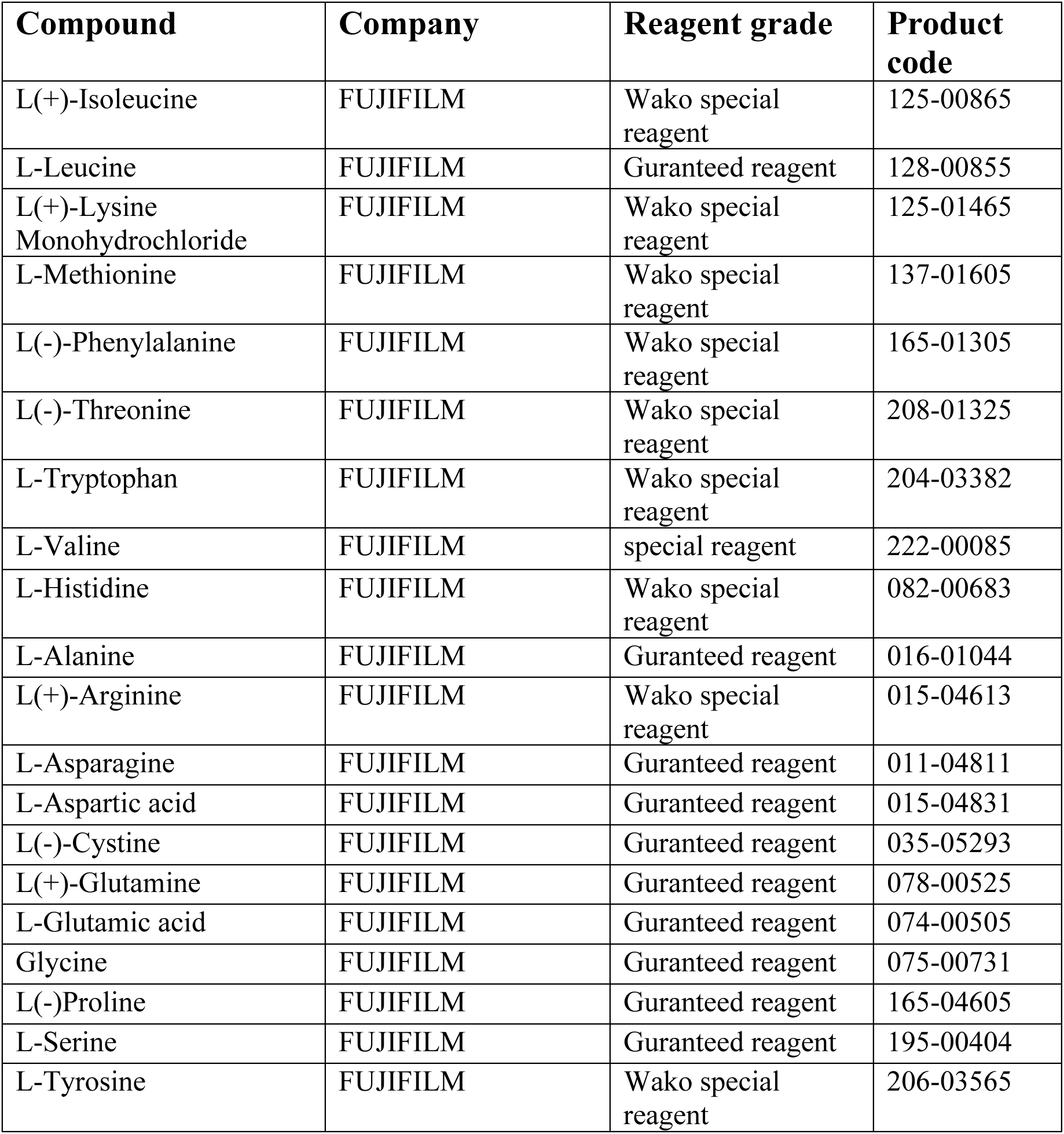
Amino acid reagents used in this study.

A week to three days before the start of the experiment, they were able to choose between the experimental control food, P30 and commercial New World primate diet (CMS-1M, CLEA Japan). The animals were completely switched to the experimental diet on day1. Subject were fed *ad libtum* with access to approximately 30g (dry matter) of food per day. In some cases, not all the diet was consumed. For example, some marmosets experienced periods without access to food, especially from nighttime to early morning, after dropping a portion out of their cage.

### Sample collection

The marmosets were fasted from 17:00 the day before. Sampling was performed the next morning. Urine samples were collected by manually expressing the bladder directly on to a clean dry tray. The blood samples were collected with heparin sodium 5,000 units/5 mL (224122458, Mochida Pharmaceutical Co. Ltd) using 26G or 0.45×13mm needle (NN-2613S, TERUMO) from the femoral veins. Restraints were used to hold the marmosets still during sampling. Plasma samples was obtained by quickly centrifuging the blood samples at 1,500 g for 10 minutes. Plasma and urine samples were stored in −80 degrees. Body weight was also measured right after sampling.

### Plasma and urine analysis

Plasma FGF21 levels were quantified using quantitative sandwich enzyme immunoassay (DF2100, FGF21 Human ELISA kit Quantikine, RSD) according to the manufacturers recommended protocol. Briefly, plasma samples were pipetted into the microplate pre-coated with human FGF21 antibody. Horse radish peroxidase was used as the detection enzyme.

Serum blood urea nitrogen (BUN), and albumin (ALB) levels were measured using FUJI DRYCHEM NX500 according to the manufacturers recommended protocol.

For quantification of amino acids, Ultra-performance LC–MS-8060 NX (Shimadzu) based on the Primary metabolites package v.2 (Shimadzu) was used to quantify plasma and urine metabolites. Ten microliter of urine or plasma samples were mixed with 150 μl of 80% methanol containing 10 μM of internal standards (methionine sulfone and 2-morpholinoethanesulfonic acid) then deproteinized by mixing 75 μl of acetonitrile. The samples were further filtered using a 10-kDa centrifugal device (Pall, OD010C35). The samples were then completely evaporated using a centrifugal concentrator (TOMY, CC-105) and resolubilized in ultrapure water. The samples were then injected to the LC–MS/MS with a PFPP column (Discovery HS F5 (2.1 mm × 150 mm, 3 μm), Sigma-Aldrich) in the column oven at 40 °C. A gradient from solvent A (0.1% formic acid, water) to solvent B (0.1% formic acid, acetonitrile) for 20 min was used to separate the samples. Multiple reaction monitoring (MRM) parameter were optimized by injecting the standard solution, then peak integration and parameter optimization were performed by a software (LabSolutions, Shimadzu). To quantify the metabolites, a standard curve drawn from serial dilutions (0.01–10 µM) were used.

### Statistics

Statistical analysis for metabolomics was performed using Metaboanalyst 6.0^54^. For normalization, the values were log transformed, and Z scores were calculated using the auto scale option. Principal component analysis (PCA) was performed and plotted.

All other statistical analysis was performed using GraphPad Prism 10. The detailed statistical tests utilized for each figure are stated in the figure legend. Briefly, repeated measures one way ANOVA was performed to assess the effects of dietary intervention. When the ANOVA testing showed significant interactions, Dunnet or Šídák post hoc testing was applied. For all statistical tests, *p*<0.05 was considered statistically significant. The *p*-values determined by post hoc testing is shown in the figures. Results of all the statistical analysis is stated with the raw data. Bar graph with overlaid lines indicating individual trajectories were generated by R, and all other plots were created by GraphPad Prism 10.

## Acknowledgement

This study was financially supported by AMED-PRIME 20gm6310011(FO), Japan Society for the Promotion of Science 22H02769 (FO), JST-FOREST JPMJFR2337 (FO), Takeda Science Foundation (FO), The Mitsubishi Foundation (FO) and by AMED under Grant Number JP23wm0625001 (TH) and JP24wm0625122 (TH). We would like to acknowledge all the caretakers (especially M.I.) of the marmoset facility in RIKEN BDR for their assistance in daily care of the animals. We also thank all the members of Obata lab, especially Chisako Sakuma and Ayano Oi for critical reading of the manuscript.

## Author contributions

F.O. conceived the project. A.U. and T.H. provided animal resources and technically assisted the animal experiment. T.I. and Y.F. performed the experiments and collected research samples. Y.F. primarily performed analysis. Y.F. and F.O. wrote the initial manuscript. A.U., T.H., and F.O. supervised the study. All authors contributed to the design of the study and reviewed the manuscript.

## Competing interests

The authors declare no competing interests.

## Data availability

All data analyzed during this study are included in this published article (and its supplementary information files). The remaining raw data are available upon request to the corresponding author.

## Supplementary Figures

**Supplementary Figure 1.**
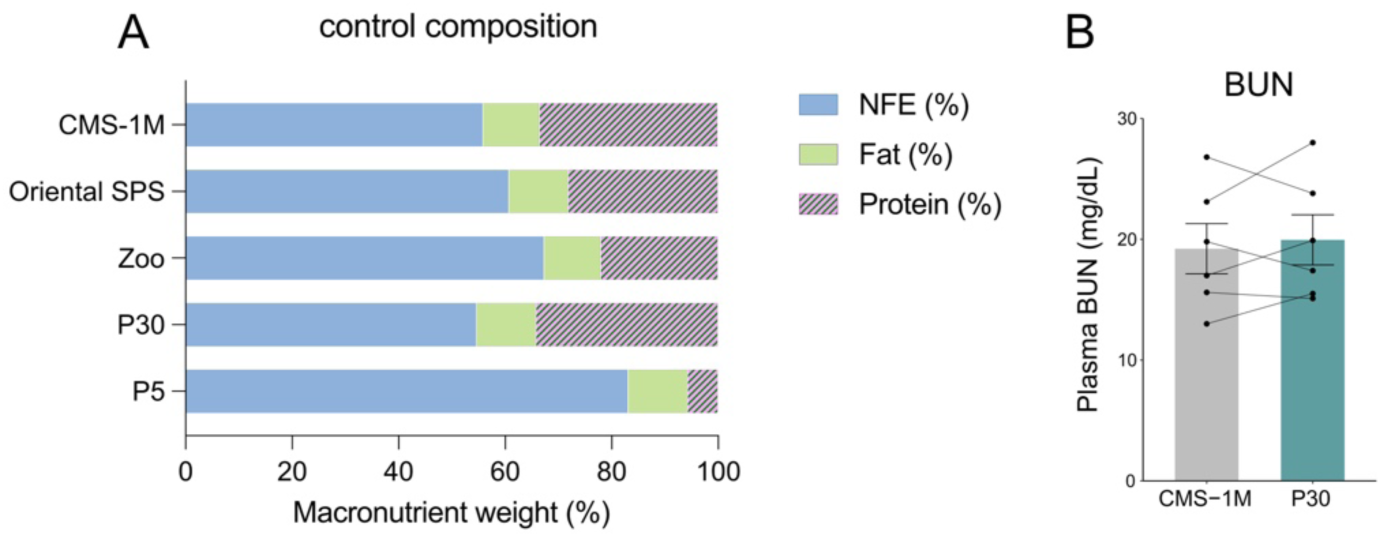
Establishing the control diet (P30) (A) Dietary macronutrient composition of commercial pellet food from SPS (Oriental yeast) and CMS-1M (CLEA Japan), survey result of a Japanese Zoo, as well as control diet (P30) and low protein diet (P5) used in this study. Nitrogen free extract (NFE), fat and protein are shown in blue, green and pink stripes, respectively. (B) Plasma blood urea nitrogen (BUN) levels of marmosets on CMS-1M then switched to P30 diet for two weeks. For (B), *p* values were determined by paired student t-test and stated on the figure. All data are presented as mean ± s.e.m.

**Supplementary Figure 2.**
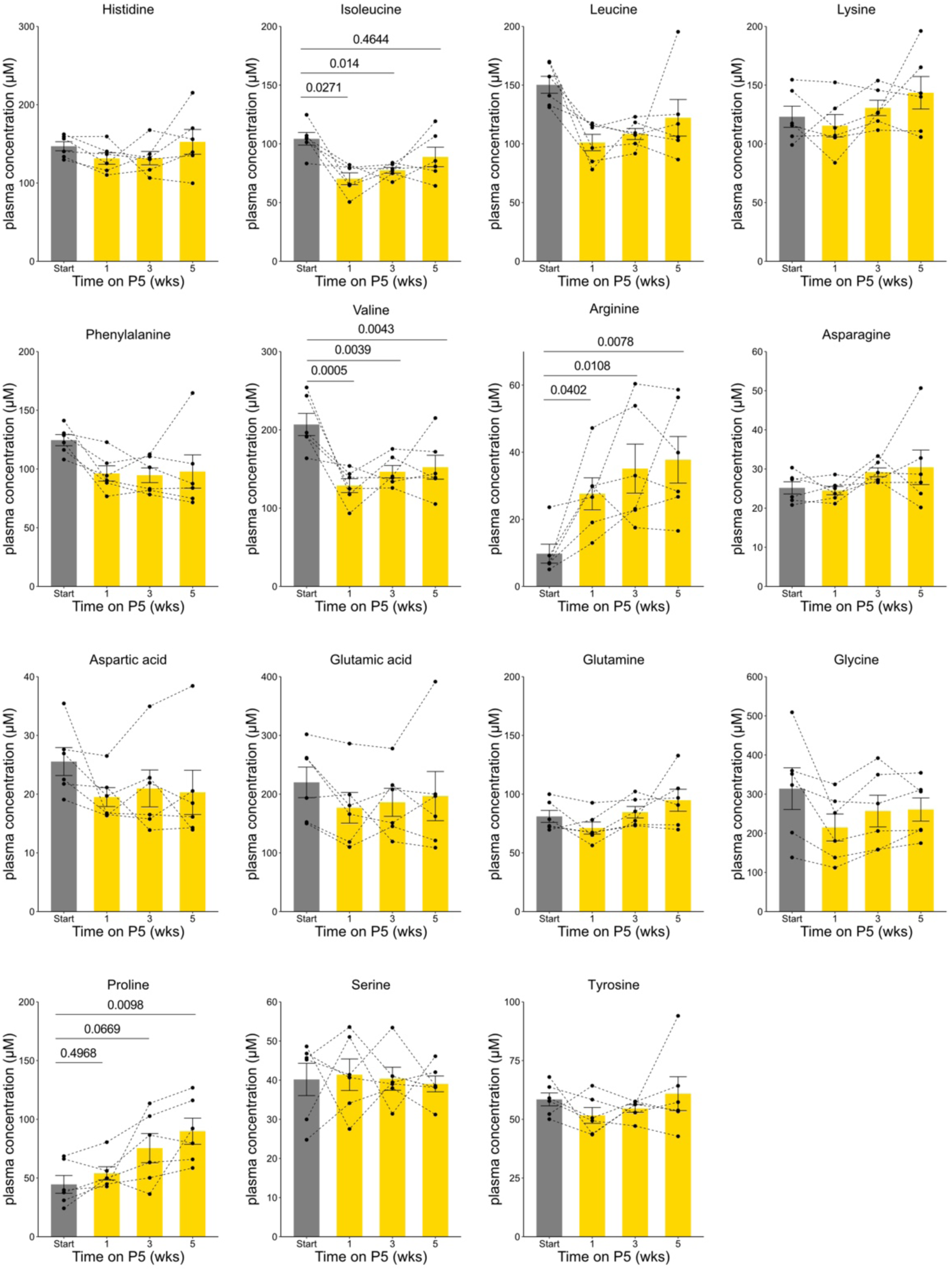
Plasma amino acid concentration under P5 diet Quantification of essential and non-essential amino acids under one to five weeks of P5 diet. Statistical analysis was performed by repeated measures one-way analysis of variance (ANOVA). The *p* values determined by post hoc analysis with Dunnet’s multiple comparison test are shown in the figure. All data are presented as mean ± s.e.m.

**Supplementary Figure 3.**
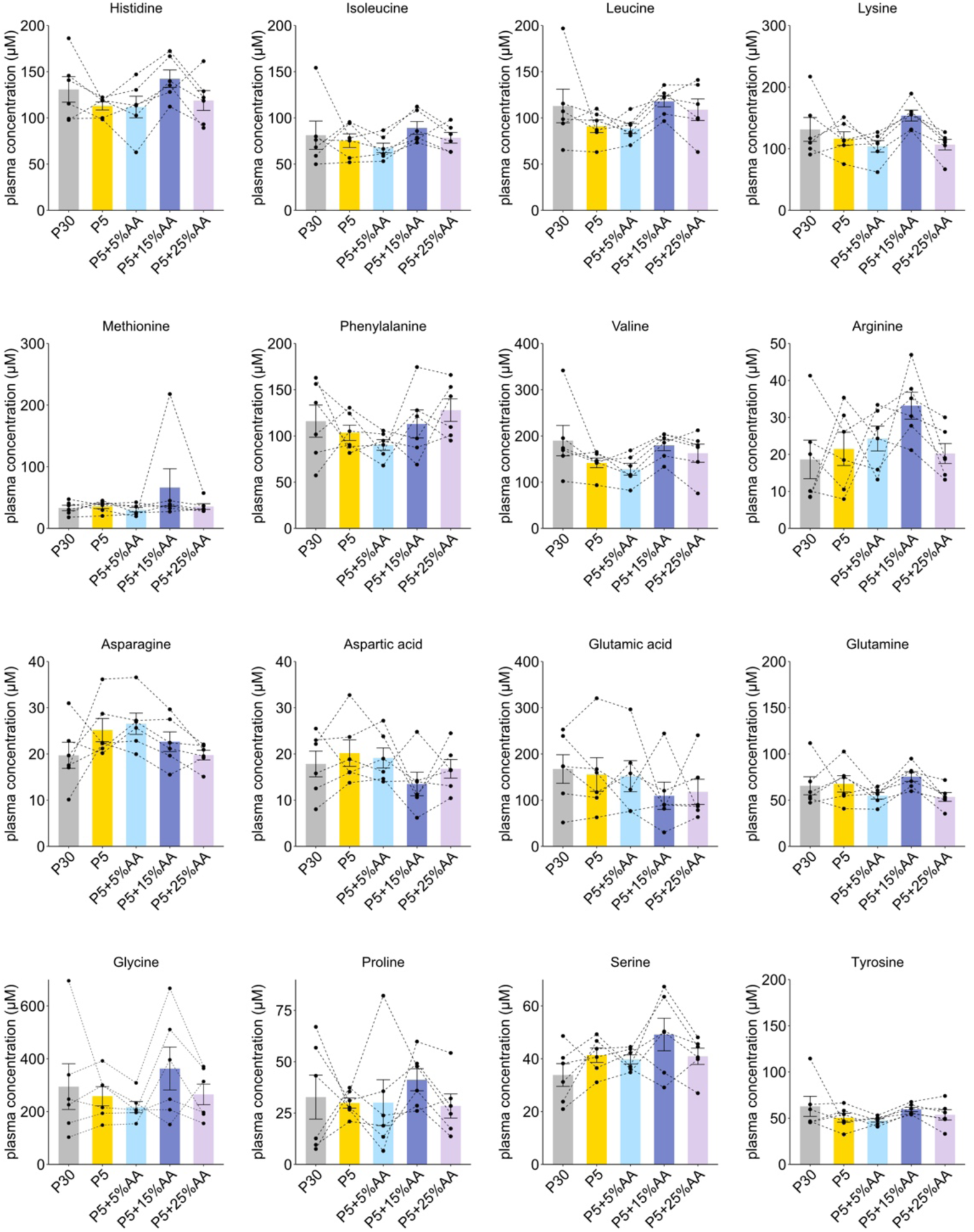
Amino acid profile under free amino acid supplementation Quantification of plasma amino acids in male marmosets. *p* values were determined by repeated measures one-way analysis of variance (ANOVA). All data are presented as mean ± s.e.m.

## Notes

### Competing Interest Statement

The authors have declared no competing interest.

